# Tetracycline transactivator overexpression in keratinocytes triggers a TRPV1 primary sensory neuron-dependent neuropathic itch

**DOI:** 10.1101/2023.07.09.548214

**Authors:** Andrew J. Crowther, Sakeen W. Kashem, Madison E. Jewell, Henry Le Chang, Mariela Rosa Casillas, Élora Midavaine, Sian Rodriguez, Joao M. Braz, Artur Kania, Allan I. Basbaum

**Affiliations:** Department of Anatomy, University of California San Francisco, San Francisco, CA 94158, USA; Department of Dermatology, University of California San Francisco, San Francisco, CA 94158, USA; Division of Dermatology, San Francisco Veteran’s Administration Healthcare System, San Francisco, CA 94121, USA; Institut de Recherches Cliniques de Montréal (IRCM), Montreal, Qc, H2W 1R7

## Abstract

Mouse models that combine tetracycline-controlled gene expression systems and conditional genetic activation can tightly regulate transgene expression in discrete cell types and tissues. However, the commonly used Tet-Off variant, tetracycline transactivator (tTA), when overexpressed and fully active, can lead to developmental lethality, disease, or more subtle behavioral phenotypes. Here we describe a profound itch phenotype in mice expressing a genetically encoded tTA that is conditionally activated within the Phox2a lineage. Phox2a; tTA mice develop intense, localized scratching and regional skin lesions that can be controlled by the tTA inhibitor, doxycycline. As gabapentin, but not morphine, relieved the scratching, we consider this phenotype to result from chronic neuropathic itch, not pain. In contrast to the Phox2a lineage, mice with tTA activated within the Phox2b lineage, which has many similar areas of recombination within the nervous system, did not recapitulate the scratching phenotype. In Phox2a-Cre mice, but not Phox2b-Cre, intense Cre-dependent reporter expression was found in skin keratinocytes which formed the area at which skin lesions developed. Most interestingly, repeated topical application of the DREADD agonist, CNO, which chronically induced G_i_ signaling in Phox2a-keratinocytes, completely reversed the localized scratching and skin lesions. Furthermore, ablation of TRPV1-expressing, primary afferent neurons reduced the scratching with a time course comparable to that produced by G_i_-DREADD inhibition. These temporal properties suggest that the neuropathic itch condition arises not only from localized keratinocyte activation of peripheral nerves but also from a persistent, gabapentin-sensitive state of central sensitization.

## Introduction

The tetracycline transactivator (tTA) system is commonly used to control transgene expression spatially and temporally in eukaryotes (Gossen and Bujard 1992). tTA protein, in the absence of tetracycline, constitutively activates transcription of downstream tetracycline response element (TRE)-containing transgenes. In this Tet-OFF configuration, exposure to tetracycline or its analog, doxycycline (Dox), arrests the expression of TRE-regulated transgenes, through inhibition of tTA binding to the TRE. In 2018, the Allen Institute introduced tTA-driven, TIGRE2.0 transgenic mouse lines as resources for neuroscientists (Daigle et al. 2018). These next-generation lines displayed increased transgene expression and reduced the complexity of mouse breeding. In most TIGRE2.0 lines, transcriptional amplification is accomplished by cloning the strong synthetic promoter, CAG, next to the tTA2 gene, in the same cassette as the downstream TRE reporter gene, which is inserted in the TIGRE locus. Owing to their impressively high expression, TIGRE2.0 lines have significantly increased the utility of cellular and activity reporters, particularly for calcium imaging of neural activity (Mohan et al. 2023).

Although the tetracycline transactivator system has been invaluable in the research of mechanism and disease processes, there have been numerous reports of tTA toxicities across cell types (Kukreja et al. 2018; Ottina et al. 2017; Jouvet et al. 2021). For example, tTA provokes hippocampal degeneration independent of known neurogenerative drivers and alters mouse behavior in some background strains (Han et al. 2012; McKinney et al. 2008). Indeed, when bred to specific Cre drivers, tTA-driven TIGRE2.0 lines can result in strain-dependent adverse events, including embryonic lethality, weight loss, cortical neuronal aberrancy, and behavioral alterations (Daigle et al. 2018). As there is minimal leak expression in these lines, the toxicity must be related to which areas are recombined and at what time. Cre drivers expressed early in embryogenesis may carry increased lethality, whereas those that are terminal fate selectors can withstand overexpression of the tTA transgene and develop normally. In certain cases, toxicity and phenotypic alterations could be prevented or reversed with doxycycline; others could not.

Phox2 is a homeodomain transcription factor highly conserved across chordates and tied to epibranchial placodes and brainstem development (Fritzsch, Elliott, and Glover 2017). In mammals, the two paralogous Phox2 genes, Phox2a and Phox2b, widely overlap within the nervous system, interact during neurogenesis, and are thought to be neural-specific (Pattyn et al. 1997; Brunet and Pattyn 2002). Within the nervous system, Phox2a and Phox2b specify developmental lineages of neurotransmitter phenotype (all noradrenergic and adrenergic neurons) and the peripheral autonomic nervous system (Tiveron, Hirsch, and Brunet 1996; Stanke et al. 1999). Of particular importance to our interest in the transmission of pain-generating signals to the brain (Wercberger and Basbaum 2019), Roome and colleagues recently reported that Phox2a is selectively expressed in progenitors that give rise to spinal cord dorsal horn projection neurons (Roome et al. 2020). Thus, with the initial goal of tracking superficial dorsal horn projection neuron activity *in vivo* through calcium imaging (Ahanonu et al. 2023), we crossed the Phox2a-Cre mouse line to selected TIGRE2.0 lines that express high levels of GCaMP6.

Unexpectedly, Cre-dependent tTA-GCaMP6 expression within the Phox2a lineage led to a remarkable, highly localized scratching phenotype. The scratching and consequent skin lesions were preventable and reversible with doxycycline. To investigate the etiology and eliminate previously described neuronal toxicity as the cause, we generated additional Cre driver crosses to various TIGRE2.0 reporter lines and searched for lesion-associated, extraneural reporter expression. Strikingly, we isolated Cre-dependent reporter expression in spatially restricted keratinocytes of the skin as the primary etiology of scratching. Chronic topical inhibition of Phox2a-keratinocytes, in which we expressed an inhibitory DREADD, promoted skin repair and arrested scratching. Furthermore, ablation of TRPV1-expressing primary afferent neurons with resiniferatoxin (RTX) reduced the frequency of scratching and led to skin repair in Phox2a; tTA-GCaMP6 mice. These results not only introduce a fascinating new mouse model of neuropathic itch but also demonstrate a chronic condition sustained by central nervous system (CNS) circuits that have been sensitized by ongoing keratinocyte-induced peripheral nerve input.

## Results

### Development of spontaneous scratching and focal skin lesions in mice with constitutive tTA-GCaMP expression in the Phox2a lineage

We crossed Phox2a-Cre mice with TIGRE2.0 lines that, after Cre-lox recombination, express tTA2, under the CAG promoter, and GCaMP6, under tTA2 control **(Figure 1A)**. These Phox2a; tTA-GCaMP mice developed significant spontaneous scratching that increased over time **(Figure 1B)** (GCaMP6s and 6f data are compiled, see Supplemental Figure 3B). The scratching and consequent development of prominent erosive skin lesions were bilateral and highly localized to the shoulder areas **(Figure 1C, Supplemental Figure 1A)**. Male and female Phox2a; tTA-GCaMP6 mice displayed similar lesions and time course of the increased scratching **(Supplemental Figure 1B)**. As nail trimming alleviated the lesions, we are confident that the erosive skin lesions were scratching-dependent **(Figure 1D)**. Interestingly, homozygosity of the tTA-GCaMP6 allele in Phox2a; tTA-GCaMP6 mice accelerated the course of skin pathology **(Supplemental Figure 1C)**.

**Figure 1:**
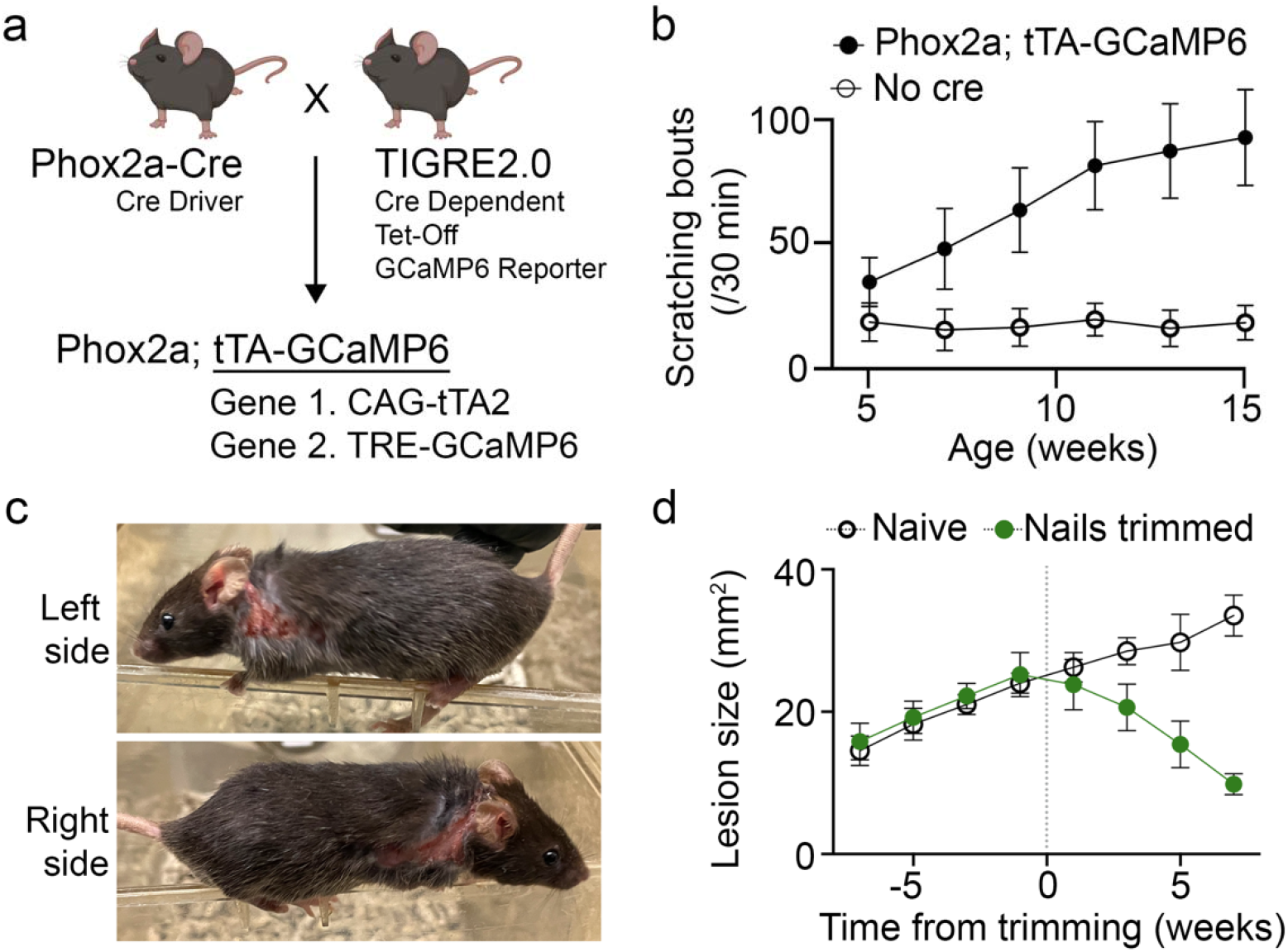
Constitutive tTA-GCaMP6 transgene expression in Phox2a+ lineage leads to localized scratching and scratching-induced skin lesions. (a) Transgenic breeding scheme to produce GCaMP6 expression off the TIGRE2.0 cassette, which encodes two genes, tTA and GCaMP6. (b) Phox2a-Cre+ mice crossed to Cre-dependent tTA-GCaMP6 mice develop spontaneous scratching, and this increases over time Cre+ (N=13), Cre-(N=15). (c) Representative images of bilateral shoulder-localized erosive skin lesions in a 12-week-old Phox2a; tTA-GCaMP6 mouse. (d) Weekly hind paw nail trimming reduces the size of skin lesions in double-transgenic mice. Trimmed nails (N=5), Untrimmed (N=4).

### The scratching phenotype in the Phox2a; tTA-GCaMP mice is the manifestation of a neuropathic itch condition

Chronic scratching in the absence of exogenous pruritogens suggests that it was a manifestation of a neuropathic itch condition that developed in the Phox2a; tTA-GCaMP6 mice. To support this categorization, we assessed pruritogen-evoked scratching and responsiveness to a first-line neuropathic itch pharmacotherapy, namely gabapentin. Injection of the potent pruritogen, chloroquine, at the lesion site, significantly elevated the frequency of spontaneous scratching above Cre-negative counterparts injected with chloroquine at the shoulder, indicating a local sensitization to pruritogens **(Figure 2A)**. Additionally, Phox2a; tTA-GCaMP6 mice, which do not have cheek lesions or increased spontaneous scratching of the cheek, showed increased scratching responses to a cheek injection of chloroquine, indicating a generalized sensitization to pruritogens **(Figure 2B).** The itch was not histaminergic, as antihistamines did not affect scratching frequency **(Supplemental Figure 2A)**. Most importantly, ip. 30 mg/kg gabapentin, significantly diminished spontaneous scratching bouts **(Figure 2C)**, consistent with our hypothesis that a neuropathic itch had developed in these mice.

**Figure 2:**
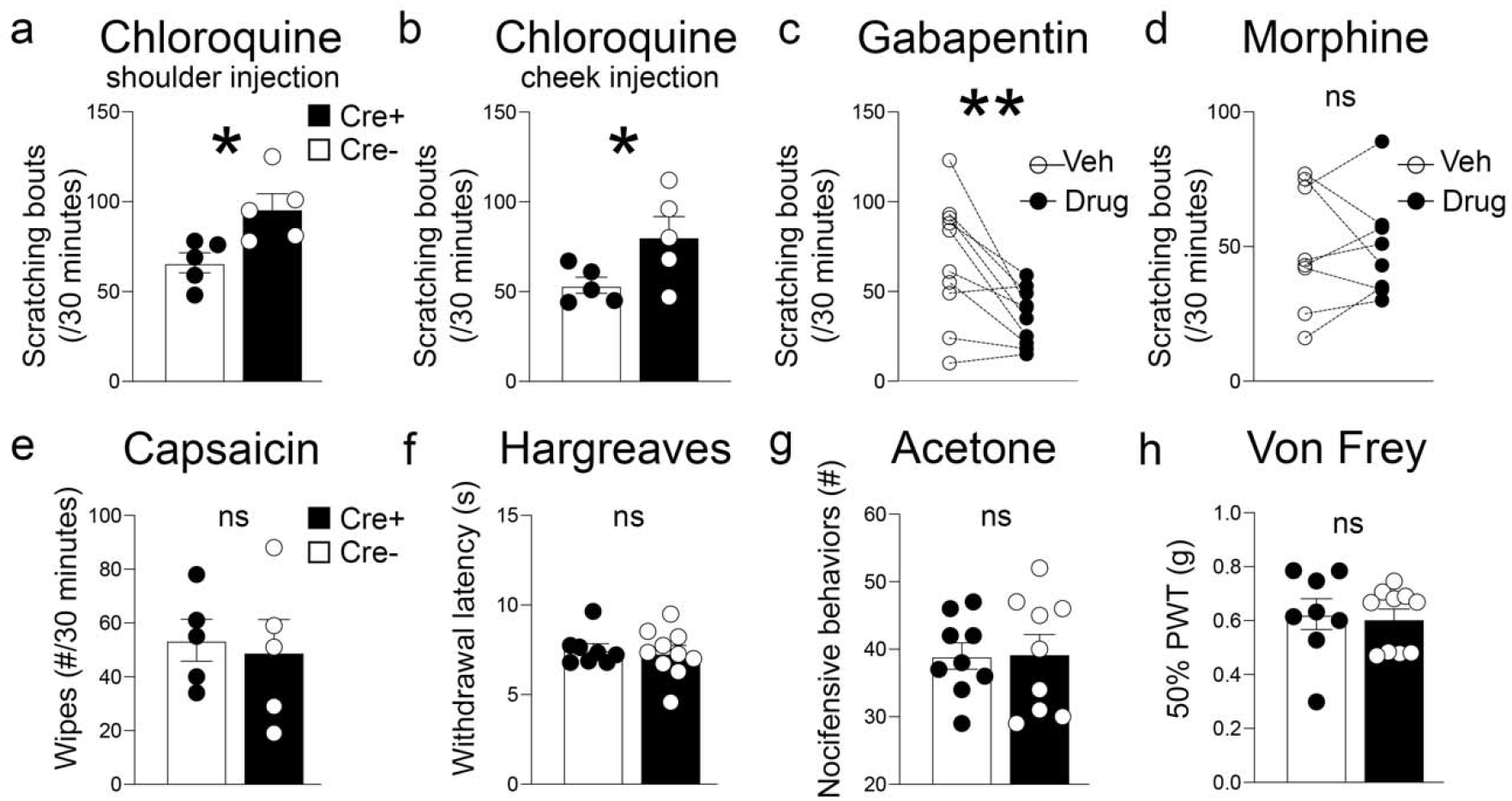
Phox2a-tTA-GCaMP6 mice develop chronic neuropathic itch, not pain. (a-b) Scratching provoked by intradermal injection of chloroquine at the shoulder (a) or at the cheek (b) in Cre+; tTA-GCaMP6 mice compared to Cre-littermate control mice. (c) Gabapentin 30 mg/kg, compared to an injection of the drug vehicle, alleviated scratching bouts in Cre+; tTA-GCaMP6 mice. Lines connect paired data points of the same mouse. (d) No difference in scratching bouts after ip. morphine 10 mg/kg in Cre+ mice. (e-h) No difference in Von Frey mechanical threshold, Hargreaves heat sensitivity, acetone noxious cold sensitivity or capsaicin-induced nocifensive behaviors (wiping) in Cre+; tTA-GCaMP6 mice compared to Cre-control mice. ns = non-significant. Statistics were analyzed by two-tailed unpaired (a-b, e-h) or paired (c-d) t-tests. * p < 0.05, ** p < 0.005, ns = non-significant.

Importantly, in contrast to gabapentin, an intraperitoneal injection of morphine, at a dose that can reverse tissue injury-induced heat hypersensitivity, did not reduce the frequency of scratching **(Figure 2D)**. Interestingly, although pruritus is a common side effect of opiate administration, particularly after intrathecal injection (Ballantyne, Loach, and Carr 1988; Nguyen et al. 2021), ip. 10 mg/kg morphine did not increase the scratching relative to the already heightened scratching in Phox2a; tTA-GCaMP6 mice. Considering the established contribution of Phox2a dorsal horn neurons to pain-associated behaviors (Roome et al. 2020), we also tested the hypothesis that these mice have a generalized alteration of nociceptive processing secondary to tissue injury in the skin. Measures of acute pain processing did not differ between Phox2a; tTA-GCaMP6 mice and their littermate Cre negative controls. Specifically, we found no differences in noxious heat sensitivity by Hargreaves testing, noxious cold hypersensitivity to acetone, or induced pain associated wiping behaviors after intradermal algogen cheek injection with capsaicin **(Figure 2E-G**). Also, von Frey fiber testing revealed no difference in mechanical thresholds, indicating that the behavior was not related to a mechanical allodynia (**Figure 2H)**.

### The neuropathic itch phenotype correlates with tTA expression

We next examined the genetic basis for this remarkably localized neuropathic itch presentation in Phox2a; tTA-GCaMP6 mice. To genetically dissect the contribution of the tTA2 gene or the GCaMP6 gene of the bicistronic TIGRE2.0 cassette, we bred Phox2a-Cre to three additional TIGRE lines that varied the configuration of the tTA2 and GCaMP6 vector **(Figure 3A)**. We first established that Phox2a-Cre crossed to the Ai96 line, which lacks tTA2 but contains CAG-driven GCaMP6s, did not develop spontaneous scratching or scratching-induced lesions **(Figure 3B)**. Next, by comparing two TIGRE2.0 lines, namely Ai162 and Ai148, which produce slow or fast variants of GCaMP6, respectively, we established that subtle changes in calcium binding kinetics of GCaMP6 are not a significant determinant of the progressive itch phenotype. **(Figure 3B, Supplemental Figure 3B)**. Importantly, an additional Cre-dependent tdTomato reporter gene (CAG-tdTomato, Ai9 line), which was the primary readout for our fate mapping studies, did not change Phox2a; tTA-GCaMP scratching levels **(Figure 3B, blue circles)**. Histology confirmed that cytoplasmic GCaMP6 was readily detectable in tdTomato+ cervical spinal cord neurons, of all three genotypes (Ai96, Ai148 and Ai162) crossed to Phox2a-Cre and Ai9 **(Supplemental Figure 3A)**. Taken together we conclude that constitutive genomic GCaMP6 expression is not sufficient to induce scratching, whereas a configuration that includes tTA2 is necessary for the phenotype.

**Figure 3:**
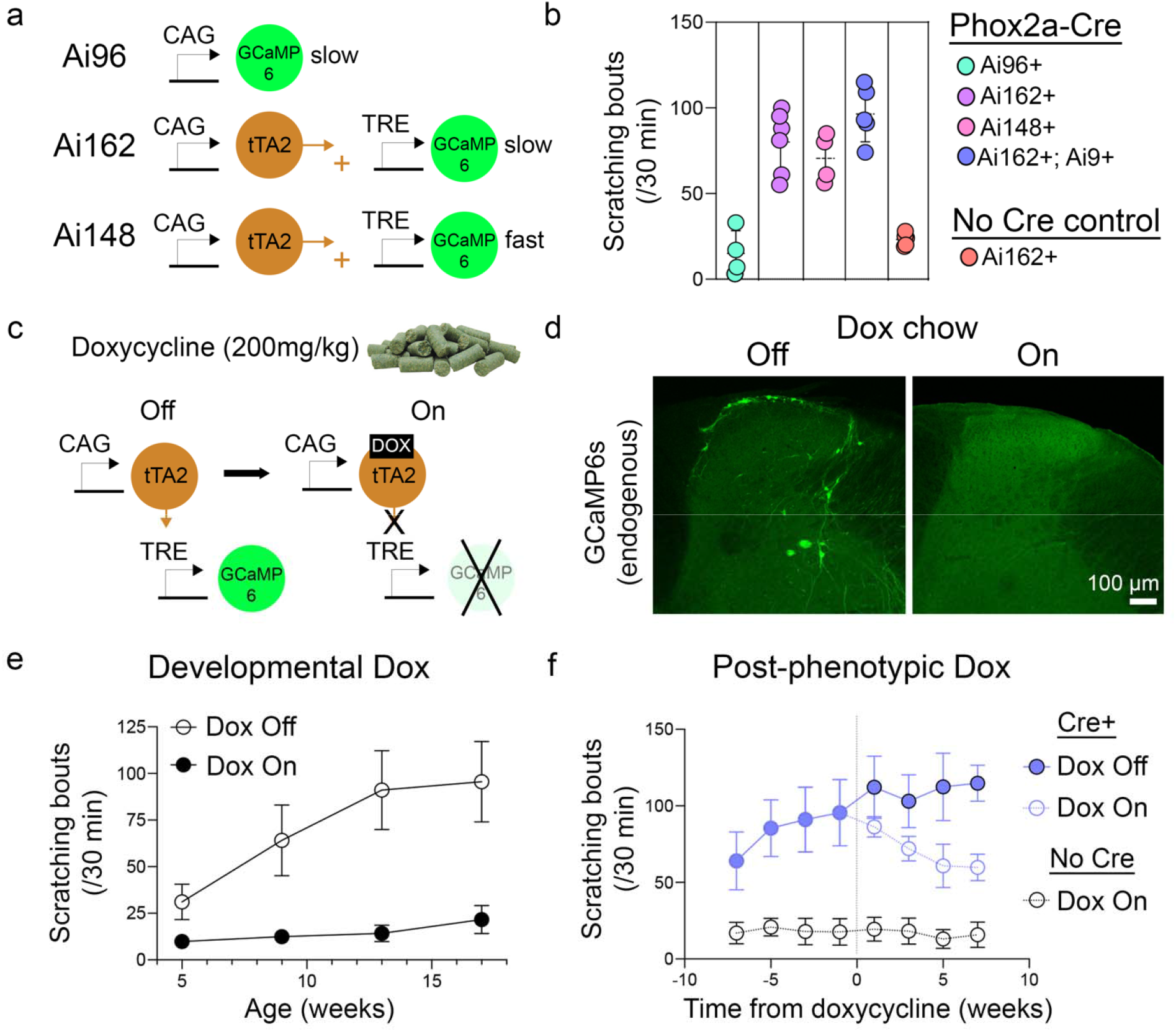
tTA inhibition reduces scratching. (a) The genetic configuration of three GCaMP6 reporter lines. The Ai96 line, without tTA transcriptional control, compared to two tTA-driven lines: Ai162 (GCaMP6s) and Ai148 (GCaMP6f). (b) Comparison of scratching frequency at 11 weeks of age across the indicated genotypes (Ai9 = Cre-dependent tdTomato reporter). Colored dots represent individual mice. (c) Doxycycline, administered through the chow, inhibits tTA activity through direct binding to the tTA protein arresting its transcriptional activity (d) Spinal cord histology confirms the downregulation of GCaMP6s in dorsal horn projection neurons of mice on Dox chow. (e) Doxycycline chow provided during pregnancy and after weaning prevents the development of scratching. Dox diet (N=5), Standard diet (N=9). (f) Doxycycline started in adulthood partially reduces spontaneous scratching. Dox On (N=4), Dox Off (N=5), Cre-(N=5).

### tTA inhibition reduces scratching in Phox2a; tTA-GCaMP6 mice

To functionally test the requirement of tTA activity for the incidence of spontaneous scratching in Phox2a-Cre; tTA-GCaMP6 mice, we administered doxycycline (Dox) through the chow (200 mg/kg). Here the Dox chronically inhibits tTA, which correspondingly represses GCaMP transcription **(Figure 3C)**. In Dox-treated adult mice, we confirmed the downregulation of GCaMP protein in Phox2a dorsal horn projection neurons **(Figure 3D)** and in the noradrenergic neurons of the locus coeruleus (data not shown). To test the effects of tTA inhibition on scratching behavior, we administered Dox chow throughout the development of the Phox2a; tTA-GCaMP6 mice, including *in utero,* by its *ad libitum* Dox feeding to the pregnant dams **(Supplemental Figure 3C)**. Strikingly, developmental Dox exposure fully prevented the spontaneous scratching and skin lesions, throughout the life of the animal **(Figure 3E)**.

Somewhat unexpectedly, these developmental Dox-treated mice, when switched off Dox chow as adults, did not develop the characteristic scratching-induced lesions. However, consistent with an ongoing effect of tTA activity after the development of scratching-induced lesions, Dox feeding to symptomatic adults significantly reduced, but did not eliminate, scratching over two months **(Figure 3F)**.

### tTA configuration without the GCaMP6 gene increases skin lesion pathology

To further interrogate the tTA-based genetic etiology, we bred Phox2a-Cre to two additional TIGRE2.0 lines, neither of which contained GCaMP6. We first examined Phox2a-Cre crossed with the TIGRE-MORF-GFP (Ai166) line (Veldman et al. 2020) **(Supplemental Figure 3D)**. In the Ai166 line, the vast majority of cells after Cre recombination express tTA, but no reporter protein, while 1-2% of cells express GFP under tTA control **(Supplemental Figure 3G)**.

Surprisingly, the combination of Ai166 with Phox2a-Cre was consistently perinatal lethal. In fact, deceased (P0) pups were regularly observed that PCR genotyped as Cre and Ai166 positive, whereas their viable littermates were Cre negative **(Supplemental Figure 3E)**. Interestingly, perinatal lethality could be prevented by embryonic doxycycline exposure **(Supplemental Figure 3E)**. Most importantly, in rescued Phox2a; Ai166 mice, the phenotype was partially recapitulated. In these mice, we observed hair loss restricted to the shoulder region and occasional spontaneous scratching directed at the shoulder **(Supplemental Figure 3F)**. These findings suggest that lethality in Phox2a-Cre; tTA mice is prevented by the GCaMP6 element of Ai162/48 lines and that tTA alone is sufficient to produce the highly localized features of the Phox2a; tTA-GCaMP6 phenotype.

We also examined an intersectional TIGRE2.0 line, Ai195, in which CAG-tTA2 is activated after FLP-FRT recombination independent of the reporter gene, lox-STOP-lox-TRE-GCaMP7s **(Supplemental Figure 3H)**. To execute conditional FLP-FRT recombination, we acquired the Hoxb8-FLPo line, in which FLPo recombinase is expressed in all tissues caudal to C7 (Bohic et al. 2023) **(Supplemental Figure 3I-J)**. Strikingly, double transgenic, Hoxb8-FLPo; Ai195 mice repeatedly licked/bit at their lower abdomen region of skin, inducing skin lesions that progressed in size over weeks, analogous to the scratching-induced lesions in the Phox2a; tTA-GCaMP6 mice **(Supplemental Figure 3K)**. Skin lesion presentation happened whether or not the mouse had a Phox2a-Cre genotype that would induce GCaMP7s expression in intersectional populations. Thus, we conclude that the CAG-tTA2 element of the TIGRE2.0 cassette is sufficient to induce profound phenotypes that include skin lesions at other locations of the body.

### tTA overexpression in spinal projection neurons does not drive scratching at the shoulder

Importantly, our genetic and Dox-based inhibition studies theoretically affect all Phox2a-lineage cells, throughout the organism, and thus cannot specify how the scratching phenotype occurs at such a restricted region of the body. Importantly, as an alteration in sensory neuron function could contribute to a localized scratching phenotype, it is significant that we observed little to no reporter expression in DRG neurons of Phox2a-Cre mice **(Supplemental Figure 4A)**. Rather, because of the restricted topography of scratching in cervical dermatomes, we also tested the hypothesis that tTA selectively perturbed the function and expression of cervical dorsal horn projection neurons of the Phox2a-Cre lineage.

First, we surveyed the morphology of fate-mapped cervical dorsal horn projection neurons, in Phox2a-Cre; Ai9 (CAG-Lox-STOP-Lox-tdTomato) mice, with or without an additional allele of tTA-GCaMP6. As expected, tdTomato+ neurons were observed in laminae I and V in control mice, without tTA-GCaMP **(Figure 4A)**. Also, in mice with tTA-GCaMP, laminae I and V tdTomato+ neurons were readily identifiable and exhibited normal morphologies **(Figure 4B)**. Furthermore, the XIth cranial motoneuron population present at cervical levels was similarly expressed in both genotypes, with and without tTA-GCaMP6, as evidenced by the clustered population of tdTomato+ motoneurons in the ventral horn and their efferents forming the XIth spinal nerve **(Figure 4A-B)**. In agreement with these observations, quantitative analysis across genotypes did not reveal differences in the number of cervical tdTomato+ neurons **(Figure 4C)**. We conclude that important developmental processes of neurogenesis, fate specification, and migration are largely uninterrupted by constitutive tTA expression in Phox2a-Cre neurons of the cervical dorsal horn.

**Figure 4:**
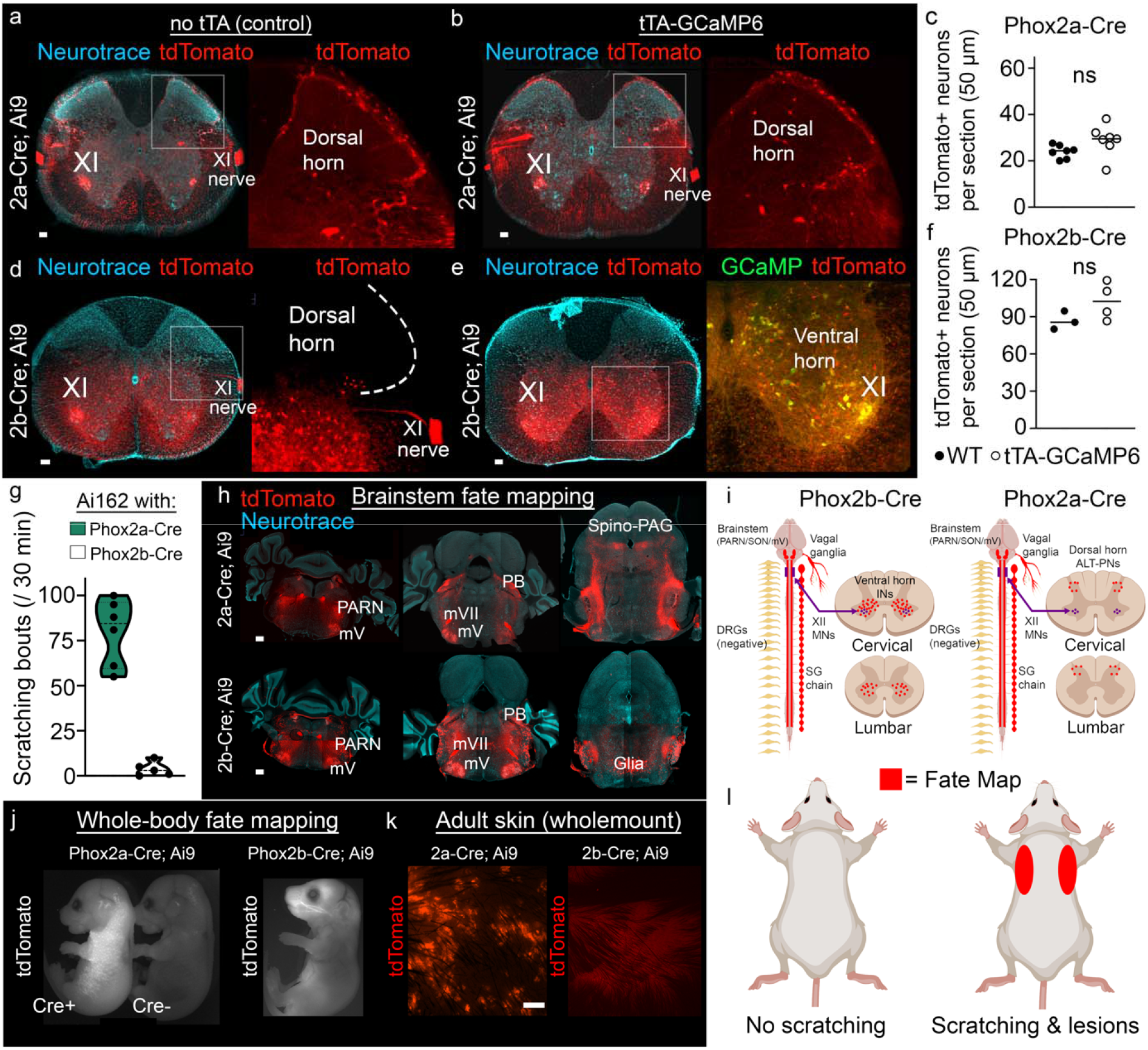
Whole-body fate mapping implicates skin expression in Phox2a-Cre mice as a determinant of the scratching phenotype. (a) At the cervical spinal cord level, Phox2a-Cre fate-mapped tdTomato+ neurons in control mice are found in their typical laminae I and V and XI motoneuron locations. (b-c) Phox2a-Cre mice with tTA-GCaMP6 show similar localization and number of labeled neurons. (d) In Phox2b-Cre; Ai9 mice the XI motoneuron pool, along with other neurons in the ventral horn, are tdTomato+, whereas the dorsal horn is devoid of labeled neurons. (e-f) GCaMP is readily apparent in many ventral horn neurons in Phox2b; tTA-GCaMP6; Ai9 mice in comparable numbers to control mice. (g) Phox2b-Cre; Ai162 mice do not recapitulate the scratching phenotype of Phox2a-Cre; Ai162 mice. (h) Comparative analysis reveals overlapping neural recombination in brainstem nuclei: mV (facial motor nucleus), PARN (parvicellular reticular nucleus) and PB (parabrachial nucleus). Spino-PAG indicates ascending terminals of dorsal horn projection neurons that are exclusively found in Phox2a-Cre mice. (i) Diagrammatic summary of fate mapping in the nervous system of Phox2b-Cre mice (left) and Phox2a-Cre mice (right). (j) Whole-body analysis illustrates tdTomato reporter expression in the skin surrounding the shoulder of Phox2a-Cre mice (left) but not in Cre-embryos (middle) or Phox2b embryos (right). (k) Phox2a lineage skin recombination was confirmed in adult skin and summarized in (l). ns = non-significant. Scale bars = 100 µm.

Next, we tested the sufficiency of tTA activity in dorsal horn projection neurons to evoke scratching. We targeted dorsal horn projection neurons in Ai162 mice using established anatomical and molecular/genetic methods. First, we injected retrograde-Cre AAV supraspinally in the parabrachial nucleus of Ai162 mice, which led to the expression of tTA in cervical spinoparabrachial projection neurons (about 50% of which are Phox2a-expressing). Second, we activated tTA in spinal NK1R-expressing neurons, which maintain chronic itch in some models of atopic dermatitis (Akiyama et al. 2015). Activating tTA by ip. tamoxifen injections in young NK1R-CreER; Ai162 mice produced cytoplasmic GCaMP6s labeling of brain and spinal cord neurons in the adult. However, even though these two approaches targeted many cervical projection neurons, neither recombination strategy provoked scratching above baseline levels over many months **(Supplemental Figure 4C)**.

### The Phox2a-Cre-, but not the Phox2b-Cre-generated fate map, includes skin keratinocytes

The above studies demonstrated that tTA-expressing projection neurons are not a sufficient driver of the scratching phenotype. However, as Phox2 genes are thought to be neural-specific, we hypothesized that other neural populations contributed to the development of this neuropathic itch phenotype. To perform a comparative analysis of the populations necessary for scratching, we bred tTA-GCaMP6 mice to a Cre line driven by the paralogous gene, Phox2b, which shares many, but not all, neuronal lineages with Phox2a. Phox2b-Cre mice crossed to tTA-GCaMP6 mice produced viable double transgenics at the expected frequency. However, these mice did not scratch **(Figure 4G)** and appeared behaviorally normal. By comparing the recombination pattern across Cre lines using Ai9 mice, we identified a shared fate map in cervical motoneurons (XI), sympathetic postganglionic neurons, brainstem motor nuclei, and the locus coeruleus **(Figure 4D-E, H-I, Supplemental Figure 4B)** and striking fate map differences in the spinal cord. Specifically, the Phox2b-Cre line only labeled ventral horn neurons **(Figure 4D-E)**. Despite the three-fold increase in the density of cervical spinal neurons in Phox2b-Cre mice compared to Phox2a-Cre, we did not detect a difference in neuronal number across tTA-GCaMP6 genotypes **(Figure 4F)**. These results, importantly, indicate again that tTA overexpression is not broadly neurotoxic.

Having ruled out several possible neural contributors to the scratching phenotype **(Figure 4I)**, we next screened for reporter expression in cells in the localized distribution of the scratching and lesions, which were selectively recombined by the Phox2a-Cre line. To screen for reporter expression outside the nervous system, we first imaged whole embryos and whole-body sections of young mice. Whole embryos revealed pronounced reporter expression in the skin of Phox2a-Cre mice. Particularly remarkable is that the expression was restricted to the shoulder region **(Figure 4J)**. Comparable reporter expression was not seen in the Phox2b-Cre mice **(Figure 4J)**. This pattern held true in the adult skin of the Phox2a and Phox2b mice **(Figure 4K)**. Moreover, cross-section histology of a 7-day-old Phox2a; Ai9 mouse revealed tdTomato fluorescence in the epidermis, and this never extended to the underlying dermis, muscle, or skin of more rostral and caudal tissue **(Supplemental Figure 4D)**.

### The topographic expression of Phox2a-Cre lineage keratinocytes matches the area of scratching-induced lesions

Strikingly, whole-embryo immunofluorescence demonstrated that tdTomato reporter expression in the Phox2a; Ai9 mice corresponded precisely with areas of focal skin lesions observed in Phox2a; tTA-GCaMP6 adults **(Figure 5A-B)**. Furthermore, using wholemount skin immunofluorescence, GCaMP signal was detected throughout adult skin lesions and colocalized with tdTomato signal in triple transgenic mice **(Figure 5B)**. Clearly, in adult mice with lesions, GCaMP expression was found in levels well above the surrounding skin or in skin lesions of IL-31 transgenic mice, which do not contain GCaMP6 (Ferda et al. 2017) **(Supplemental Figure 5A, left).** In contrast to large aggregations of GCaMP+ cells found in untreated Phox2a; tTA-

**Figure 5:**
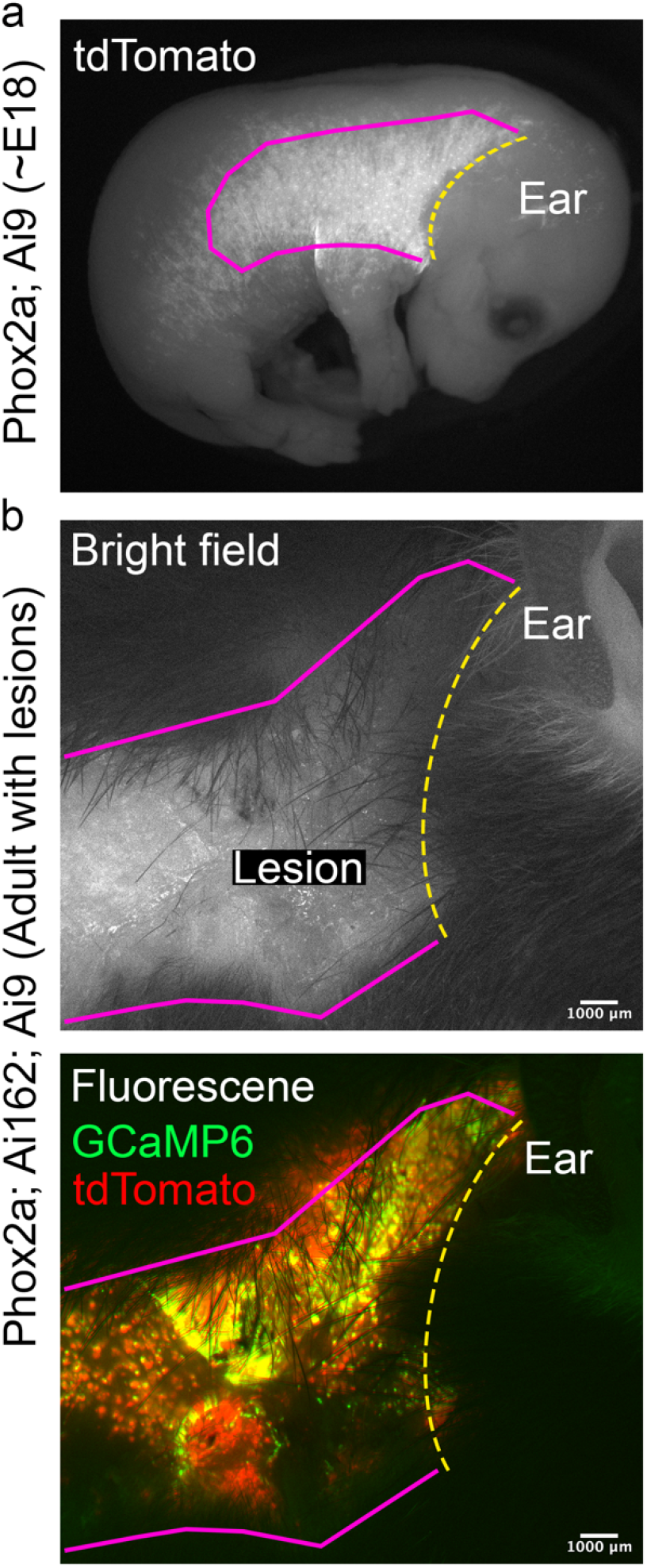
Phox2a-keratinocytes are topographically restricted to the area that matches the region of skin lesion development. (a) Visualized by tdTomato pseudo-colored in white, a Phox2a; Ai9 embryo reveals a striking pattern outlined in pink and yellow dotted line, which defines the neck boundary of recombination. (b) The hairless area of skin seen in the bright field image (top) indicates the extent of a skin lesion in a Phox2a; Ai9; Ai162 mouse. The area of skin lesion below the yellow dotted line mirrors the pink outline of reporter expression topography in (a). Under fluorescent imaging (bottom), green (GCaMP6) and red (tdTomato) signal fills the entire lesioned area, but not the surrounding skin.

GCaMP6 mice, mice with developmental doxycycline treatment presented with GCaMP signal in a normal appearing keratinocyte distribution **(Supplemental Figure 5A, top right)**. Additional images of embryos in Ai166 mice reveal single keratinocytes amongst the entire recombined lineage and prominently show full epidermal coverage and hair follicle structures **(Supplemental Figure 5B)**. Suggesting that Phox2a-keratinocytes exert a pathophysiological effect by interacting with the peripheral sensory neurons, we observed free-ending sensory nerve fibers near tdTomato+ keratinocytes of the epidermis and hair follicles **(Supplemental Figure 5C)**.

### Keratinocyte signaling is required for the maintenance of spontaneous scratching and skin lesions in Phox2a; tTA-GCaMP6 mice

To test the contribution of keratinocytes to the scratching phenotype, we inhibited keratinocytes by activation of G*_i_* signaling in Phox2a; tTA-GCaMP6 mice that also contained a Cre-dependent hM4Di inhibitory DREADD in their genome **(Figure 6A)**. We applied CNO, or phosphate-buffered saline on the side contralateral to CNO, by topically painting liquid on the skin and lesions every day for 10 days **(Figure 6B)**. **Figures 6C-E** show that chronic topical application in the Phox2a; tTA-GCaMP6; hM4Di mice significantly reduced scratching bouts and reversed skin lesions, but only on the side of CNO application. Somewhat unexpectedly, topical application of CNO to the shoulder region in Phox2a-Cre; floxed-hM3Dq mice, which we hypothesized would activate local keratinocytes, did not elicit scratching. We conclude that ectopic keratinocyte activity develops in Phox2a; tTA-GCaMP6 mice and is a necessary contributor to this novel neuropathic itch condition.

**Figure 6:**
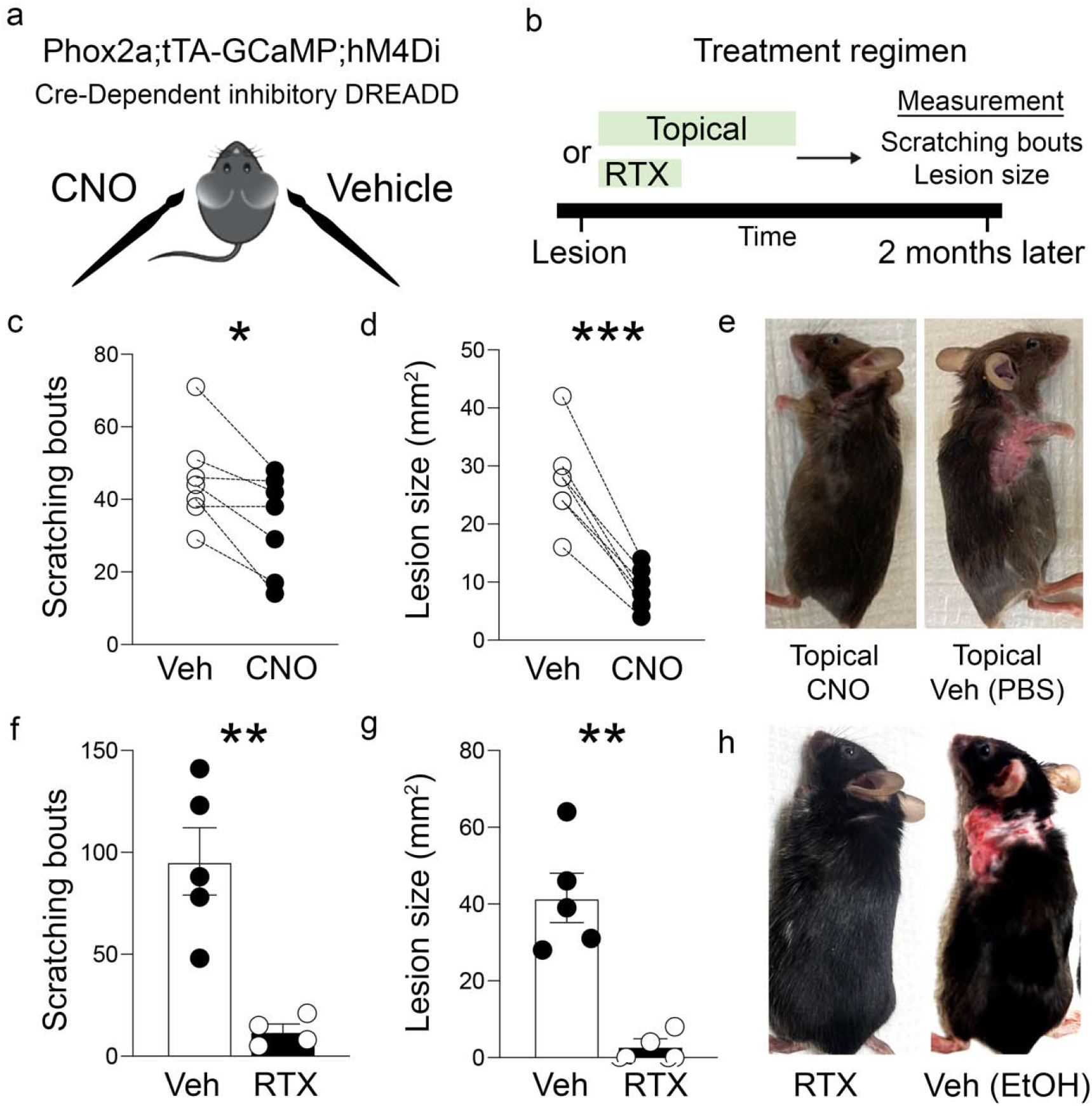
TRPV1-expressing primary afferent neurons contribute to the neuropathic itch condition in Phox2a; tTA-GCaMP6 mice. (a) Topical and unilateral administration scheme to modulate keratinocyte function using inhibitory DREADDs. (b) CNO was topically applied repeatedly to phenotypic mice over 10 days, and scratching and lesion size were assessed after 2 months. In a separate experiment, resiniferatoxin was given to phenotypic mice in three successive doses, and scratching and lesion size were assessed 2 months after the last injection. (c-e) CNO-stimulated Gi-coupled signaling reduces scratching and skin lesions. (f-h) Ablation of TRPV1+ nociceptors with resiniferatoxin reduces scratching and skin lesions in transgenic mice. Statistics were analyzed by two-tailed paired (c-d) or unpaired (f-g) t-tests. * p < 0.05, ** p < 0.005, *** p < 0.0005.

### TRPV1-expressing primary sensory neurons contribute to the scratching and skin lesions in Phox2a; tTA-GCaMP6 mice

We next addressed the mechanism through which keratinocytes communicate with itch-provoking circuits in the nervous system. As the majority of pruritogens initiate scratching by activating TRPV1-expressing afferents, we ablated TRPV1+ sensory neurons with the capsaicin super-agonist resiniferatoxin (RTX) and tracked scratching and lesion size after the ablation **(Figure 6B)**. Consistent with the ablation of the TRPV1 population, these mice demonstrated significantly increased latency to noxious heat compared to controls, but not to mechanical stimulation (data not shown). Most importantly, **Figures 6F-H** show that spontaneous scratching and skin lesions were completely reversed at 2 months post-ablation. Even though ablation of the TRPV1 nociceptors and pruritoceptors occurs within days of RTX injection, it is of interest that the scratching only declined after several weeks, with additional time needed for the skin lesions to disappear. Based on this prolonged time course, we conclude that the keratinocytes communicate with these TRPV1-expressing afferents to drive the neuropathic itch phenotype, but that induced and persistent hyperexcitability of dorsal horn circuits contributes to and can sustain the phenotype for an extended period.

## Discussion

This study introduces a neuropathic itch model that develops spontaneously in Cre-lox-based double-transgenic mice and is characterized by a remarkably confined distribution of the scratching and consequent skin lesions. This focal disorder stems primarily from the selectivity of the Phox2a-Cre lineage, where conditional tTA activation in shoulder keratinocytes specifically triggers scratching at their location. That conclusion is supported by the selective inhibition of Phox2a-keratinocyte activity using topical DREADD-based methods, which reversed the localized skin lesions and normalized scratching. Furthermore, as resiniferatoxin and gabapentin alleviated the scratching phenotype, we conclude that TRPV1-expressing primary afferent neurons and dorsal horn circuitry downstream of the keratinocytes are necessary contributors to the development of neuropathic itch in this model.

### Temporal properties of the neuropathic itch model

It is of particular interest that only after a significant two-month delay did ablation of the TRPV1 subset of primary sensory neurons reverse the scratching phenotype. Moreover, a delay in the alleviation of scratching could also be induced in conditions where TRPV1 sensory neurons were intact. Specifically, although DREADD-mediated inhibition of Phox2a-keratinocyte activity did reduce the scratching phenotype, this only occurred after repeated dosing of CNO, on a similarly extended timescale to that induced by resiniferatoxin injection. These observations are also relevant to our finding that inhibiting the primary genetic cause, tTA, by the administration of doxycycline, only partially relieved the localized scratching after the development of lesions. Taken together, we conclude that while this neuropathic itch model is initiated in the periphery, it can be sustained, although not permanently, by sensitized circuits in the nervous system. Consistent with our conclusion that a CNS-localized central sensitization process underlies the persistence of the neuropathic itch model, gabapentin, which exclusively acts on CNS circuitry, reduced the scratching.

### Keratinocyte-derived mechanisms that produce itch

Scratching is well-known to be provoked by mediators released by epidermal cells and keratinocytes, in particular, have been shown to drive itch through direct communication to primary sensory neurons in multiple contexts (Ikoma et al. 2006). Our group, as well as others, have demonstrated that mice in which protease activated receptor 2 is overexpressed in keratinocytes develop hypersensitivity to pruritogens and house dust mites (Frateschi et al. 2011; Braz et al. 2021). Constitutive expression of Ca++ activated KCa3.1 channel in keratinocytes also leads to intense pruritus and skin lesions (Lozano-Gerona et al. 2020). And under inflammatory and dry skin contexts, keratinocytes can release cytokines, including thymic stromal lymphopoietin (TSLP) and IL-33, which can directly activate sensory nerves to trigger pruritus (Wilson et al. 2013; Trier et al. 2022; Liu et al. 2016). Interestingly, optogenetic activation of keratinocytes implicated calcium signaling in keratinocytes as a TRPA1-dependent initiator of TSLP secretion and evoked pruritus (Wilson et al. 2013).

### Spatial properties of the scratching phenotype

Regarding the intriguing localization of the phenotype, it is unclear when the Phox2a gene, and thus Cre, is transcriptionally activated in keratinocyte precursors. Skin expression is surprising considering the commonly held view that Phox2 genes are neural-specific. Is recombination attributable to patterned expression in the ectoderm or in terminally differentiated keratinocytes? Also, could the phenotype be a consequence of BAC-based genetic engineering? In a previous study using a BAC-form of Phox2b-Cre, only preganglionic cholinergic parasympathetic neurons were labeled, which is different from what was predicted from molecular expression studies and knock-in Cre mice (Rossi et al. 2011; Hirsch et al. 2013). Importantly, minimizing the concern of non-physiological recombination in the BAC-derived Phox2a-Cre mice used in this study, skin reporter expression is also found in a knock-in Phox2a-Cre (Kania et al, unpublished data). Perhaps the remarkable localized phenotype is associated with the contribution of Phox2a to orchestrating the special instance of accessory nerve exit and the development of branchial structures dating back to early vertebrates (Fritzsch, Elliott, and Glover 2017).

### Mechanisms of toxicity

Consistent with the initial TIGRE 2.0 report (Daigle et al. 2018), we confirm that expression of the CAG-driven tTA2 allele during development is the determining factor for pathology. Itch-associated lesions were found only in mice with active tTA during critical developmental periods, and an exaggerated pathology was observed in tTA-GCaMP6 homozygous mice, indicating a strong genetic basis. Interestingly, studies of tTA in various organisms suggest that high levels of expression can induce cellular dysfunction, including non-specific transcription events, aberrant protein ubiquitination, or transcriptional squelching (Gong et al. 2005; Gill and Ptashne 1988; Berger et al. 1990; Damke et al. 1995; Salghetti et al. 2001). Other studies reported that TIGRE1.0 TRE-GCaMP mice that express CaMKIIa-tTA exhibit varying cerebral ictal events depending on the GCaMP variant (Steinmetz et al. 2017). In contrast, in our Phox2a-Cre model, the GCaMP6 variant did not significantly impact the scratching phenotype, and typical GCaMP toxicity signatures, for example, ectopic nuclear localization associated with neuronal dysfunction, were not observed (Yang et al. 2018). Even more paradoxically, replacing the GCaMP6 gene with GFP in the Ai166 mice resulted in perinatal lethality. Our interpretation of this result is that GCaMP6s expression causes slight epigenetic suppression of the locus and lower tTA levels.

### Future experiments

Based on our findings, we anticipate that selectively expressing tTA in keratinocytes using a keratinocyte Cre driver in Ai162 mice will lead to the development of scratching-induced lesions that are not limited to the shoulder area. Notably, our experiments in which CAG-tTA2 is activated in multiple tissues (neurons, muscle, skin) primarily exhibited progressive skin issues. This observation suggests that tTA selectively evokes pathology within skin tissue. Conducting transcriptomic analysis of keratinocytes that express tTA and have a profound impact on the propensity for itch in mice may yield valuable insights into the downstream mediators that engage the peripheral nervous system. Further transcriptomic analysis of tTA-expressing skin tissue may also provide valuable insights into the keratinocyte signaling modulated by G_i_-DREADD coupled activation. Previously, activation of G_i_ signaling in keratinocytes using a K5-Cre; LSL-hM4Di mice treated with CNO led to proliferation and prevention of differentiation (Pedro, Parra, and Iglesias-Bartolome 2019). It is possible that G_i_ signaling in the context of transgene-evoked keratinocyte pathology allows for the restoration of epidermal integrity through alteration of basal keratinocyte differentiation or repair of the skin ulceration. Clearly, it will be important to discern the mechanism whereby G_i_ signaling counteracted the tTA effects in keratinocytes and restored skin integrity and reduced itch. Finally, as keratinocytes of the glabrous skin have been implicated in mechanical pain hypersensitivity, it will be important to identify the contextual and topographical determinants that influence keratinocytes to trigger pain or itch (Baumbauer et al. 2015; Moehring et al. 2018; Mikesell et al. 2022).

### Conclusion

This study describes a profound scratching phenotype driven by molecular and topographical factors, highlighting the dysregulation of keratinocytes, nervous system responsivity, and the provocation of itch. Our finding underscores the importance of careful selection of appropriate transgenic mouse lines and consideration of transgene toxicities. Importantly, when planning to use the TIGRE2.0 lines, we recommend using inducible recombination strategies, if possible, such as CreER lines or viral approaches in adult mice, to avoid developmental expression of the CAG-tTA2. Lastly, our ability to completely reverse skin lesions through topical activation of G_i_ signaling in keratinocytes may have implications for other skin conditions that rely on extracellular communication with the peripheral nervous system.

## Methods

### Transgenic mouse breeding and maintenance

Animal husbandry and care were provided by the UCSF LARC. Animal procedures were conducted in accordance with the guide for the Care and Use of Laboratory Animals, as adopted by the NIH, and with approval of the Institutional Animal Care and Use Committee at the UCSF and performed in facilities accredited by the Association for the Assessment and Accreditation of Laboratory Animal Care International (AAALAC). Phox2a-Cre mice were provided by Artur Kania (Université de Montréal) (Roome et al. 2020). Ai162 (#:031562), Ai96 (#:028866), Ai148 (#:030328), Ai166 (#:035404), Ai9 (#:007909), floxed-hM4Di (#:026219), floxed-hM3Dq (#:026220) and Phox2b-Cre (#:016223) mice were obtained from the Jackson Laboratory. NK1R-CreER mice were provided by the Ross lab (Huang et al. 2016). Both male and female mice were studied. All JAX-derived mice were maintained on the C57BL/6 background or backcrossed at UCSF to C57BL/6 background for at least 3 generations. Animals were separated by sex at weaning and group housed at 3-5 animals per cage. Only virgin animals were used for experiments. All mice were fed a standard diet unless otherwise noted (to receive doxycycline) in standard cages and were maintained on a 12-hour light/dark cycle.

### Doxycycline administration

Doxycycline chow 200 mg/kg was purchased from Bio-Serv (S3888) and stored at 4 degrees. Dox chow was offered to mice *ad libitum* and the chow was refreshed with Dox chow from storage every 2 weeks. To breeding females, the doxycycline chow began before mating and continued through to weaning. For post-phenotypic Dox, mice that never received Dox, but displayed lesions, were administered Dox ad libitum for 2 months when lesions were >20 mm^2^.

### Measurement of mouse behavior

For all mouse behavior experiments, the experimenter was blinded to the extent possible. Mice were acclimatized to the behavior room and habituated for 3 straight days for 30 minutes to the experimental apparatus.

#### Documenting scratching

Mice were acclimatized as above and placed in opaque white cylinders over a platform and recorded via on ImagingSource camera for 15 or 30 minutes in the dark. In some settings, mice were recorded with a handheld Sony video camera on a mirrored platform for a similar duration. Scratching bouts were scored on video playback and were defined as a single hind paw motion that contacts either the cheek or back. The end of a bout was defined as when the hind paw was lowered.

#### Chloroquine-induced scratching

Mice were acclimatized for 30 minutes, and then 200 μg of chloroquine (Sigma 1650000) was injected intradermally into the cheek or intra-lesional at the shoulder or in a comparable spot in a Cre-mouse without lesions. Scratching was measured for 30 minutes, as described above.

#### Capsaicin-induced nocifensive behaviors

Mice were acclimatized for 30 minutes, and then 20 μg of capsaicin (Sigma M2028) was injected intradermally in 20 μL of 7% Tween-80, 10% ethanol in phosphate-buffered saline (PBS) into the defined anatomical sites of the cheek. Wiping was measured for 30 minutes in a setting comparable to that used to analyze scratching (Shimada and LaMotte 2008).

#### Von Frey measurement of mechanical sensitivity

For all groups, we recorded 3 days of baseline mechanical sensitivity. Animals were habituated on a wire mesh for 2 hours, after which we used von Frey filaments (sizes 1.65, 2.44, 2.84, 3.22, 3.61, 3.84, 4.08, and 4.31) to measure mechanical withdrawal thresholds, using the up–down method. These filaments correspond to the following weights: 0.008, 0.004, 0.07, 0.16, 0.4, 0.6, 1, and 2 g, respectively.

#### Hargreaves measurement of heat sensitivity

We acclimatized the mice for 30 min in Plexiglass cylinders. The mice were then placed on the glass of a Hargreaves apparatus, and the latency to withdraw the paw from the heat source was recorded. Each paw was tested five times, and we averaged latencies over the five trials. Hargreaves tests were done 1 hour after the von Frey measurement.

#### Acetone-induced measurement of cold sensitivity

Mice were habituated for 30 min on a mesh in plexiglass cylinders. Next, we used a syringe to squirt 50 µl acetone onto the plantar surface of the paw. The responses of the mice directly after the application of acetone were recorded on video for 30 s. Each paw was tested five times and we measured events spent lifting, licking, or flinching the paw. Results are displayed as the total events across the five trials.

#### Assessment of skin lesion size

Skin lesions were defined as erosive (loss of epidermis) lesions on the back of unshaved mice. Maximal width and length of the lesions were measured with a digital caliper (Rexbeti).

### Chemical injections

*Gabapentin (Sigma G154)* was injected intraperitoneally at 30 mg/kg after 30 minutes of acclimatization. Mouse behavior was measured 30 minutes after injection for another 30 minutes.

*Morphine* (Sigma M8777) was injected intraperitoneally at 10 mg/kg after 30 minutes of acclimatization. Mouse behavior was measured 30 minutes after injection for another 30 minutes.

*Cetrizine* (Sigma BP837) was injected intraperitoneally at 30 mg/kg after 30 minutes of acclimatization. Mouse behavior was measured 30 minutes after injection for another 30 minutes.

*Resiniferatoxin* (Adipogen AG-CN2-0534-MC01) was diluted at 1.0 mg/ml in 100% ethanol and then subcutaneously injected with 3 escalating consecutive daily doses of 30, 60, and 100 μg/kg in 300 μL of normal saline. The efficacy of RTX-mediated TRPV1+ sensory nerve ablation was assessed by monitoring tail flick latency.

*Clozapine-N-oxide* (company) was injected at 1.0 mg/kg in normal saline intraperitoneally 5 minutes before behavioral testing. Alternatively, 5.0 μg in 40 μL of PBS was topically painted on the skin every day for 10 days, unilaterally over the skin lesion; the contralateral skin lesion received a similar volume of PBS.

*Tamoxifen* (Sigma-Aldrich T5648) was dissolved in Mazola corn oil (1.0 mg / 50 μl) at 55 degrees for 15 minutes with vortexing, and aliquots were stored at −20 degrees. At 4 weeks of age, mice received an intraperitoneal injection of Tamoxifen (80 mg/kg) 3 times in a 10-day period, which effectively induced NK1R-CreER recombination. Mice were returned to their home cage after injection, and their general health was monitored daily.

### Histology

For histology of whole mount samples (DRGs, spinal cord, skin, and embryos), mice were transcardially perfused with 4% formalin, the spinal column and a section of the skin of the upper back were dissected and stored in 4% formalin overnight. After washing with PBS, DRGs and the spinal cord were manually extracted from the spinal column under a dissecting microscope. In a few cases, a pregnant dam was perfused with 4% formalin near term to obtain embryos for whole-body tissue analysis, and the embryos were extracted and post-fixed in 4% formalin. For histology of sectioned samples, the spinal cord or skin was sectioned using a freezing microtome or cryostat at 50-micron thickness. Sections were stored in PBS after sectioning. Beta3-tubulin antibody was the single exception of antibody labeling used throughout the paper and was performed following standard immunohistochemistry procedures. Some DRG and spinal cord sections were stained with Neurotrace (N21479, ThermoFisher) or DAPI (D1306, ThermoFisher). Sections were mounted onto glass slides using Fluoromount mounting media (00-4958-02, ThermoFisher) and a no.1.5 glass coverslip. Various images of the endogenous expression of tdTomato or GFP (including GCaMP) were acquired using LED illumination, and Standard Detectors on an FV3000 (4, 10, or 20X objectives) or Axiozoom V16 outfitted with a 1X objective and an axioCam 712. In the sample from a rescued Phox2a; Ai166 mouse, the spinal cord and brain were cleared using Life Canvas technologies SHIELD clearing reagents and were imaged on a light-sheet microscope following protocols established by (Park et al. 2018).

For histological counts of cervical spinal neurons, 50-micron sections of the entire cervical enlargement were collected using a cryostat. A regularly spaced subset of sections (containing a section every 150 microns) was selected for DAPI staining and mounted. All selected sections were imaged for endogenous tdTomato and DAPI at 1.6 x 1.6 x 8-micron resolution with eight z-slices using a 10X objective on an FV3000 confocal. Cell counting was done on at least three sections per mouse and averaged. An unpaired t-test between groups was performed in Prism. For a cell to be counted, it had to meet the criteria of having a rounded cell body represented in the z-stack and DAPI-labeled nucleus. Cell counting was performed blind to the independent variable (tTA genotype) using the Cell Counter plugin in ImageJ (FIJI).

### Data presentation

Diagram schematics were created with BioRender.com. Graphs were generated in GraphPad Prism9. Figures were compiled in Adobe Photoshop. Statistical tests were performed in Prism9. Statistical analysis of differences between groups was performed using Student’s t-test (unpaired for comparison between two groups or paired for treatments performed in the same mouse). Data were considered significantly different at P of < 0.05. Data in bar graphs are expressed as means ± SE.

## Author Contributions

S.K., A.C., and A.B. designed the project and wrote the manuscript; A.C., S.K., M.J., and H.C. performed histology, imaging, image processing, and data analysis; S.K., M.J., and S.R. performed the behavioral testing and analysis. M.RC. performed the intracranial viral injections. M.J. assisted with manuscript preparation. E.M. and J.B. assisted with experiments.

## Acknowledgments

This work was supported by NIH NSR35097306 (A.B.), Open Philanthropy (A.B.), NIH F32 5F32DE029384 (A.C.), Canadian Institutes of Health Research (PJT-162225, MOP-77556, PJT-153053, and PJT-159839) (A.K.), NSF Graduate Research Fellowship 2034836 (M.R.C.), and a Dermatology Foundation Career Development Award, Investigator Research Fellowship and NIH-5T32AR007175 (S.K.).

---

**Supplemental Figure 1:**
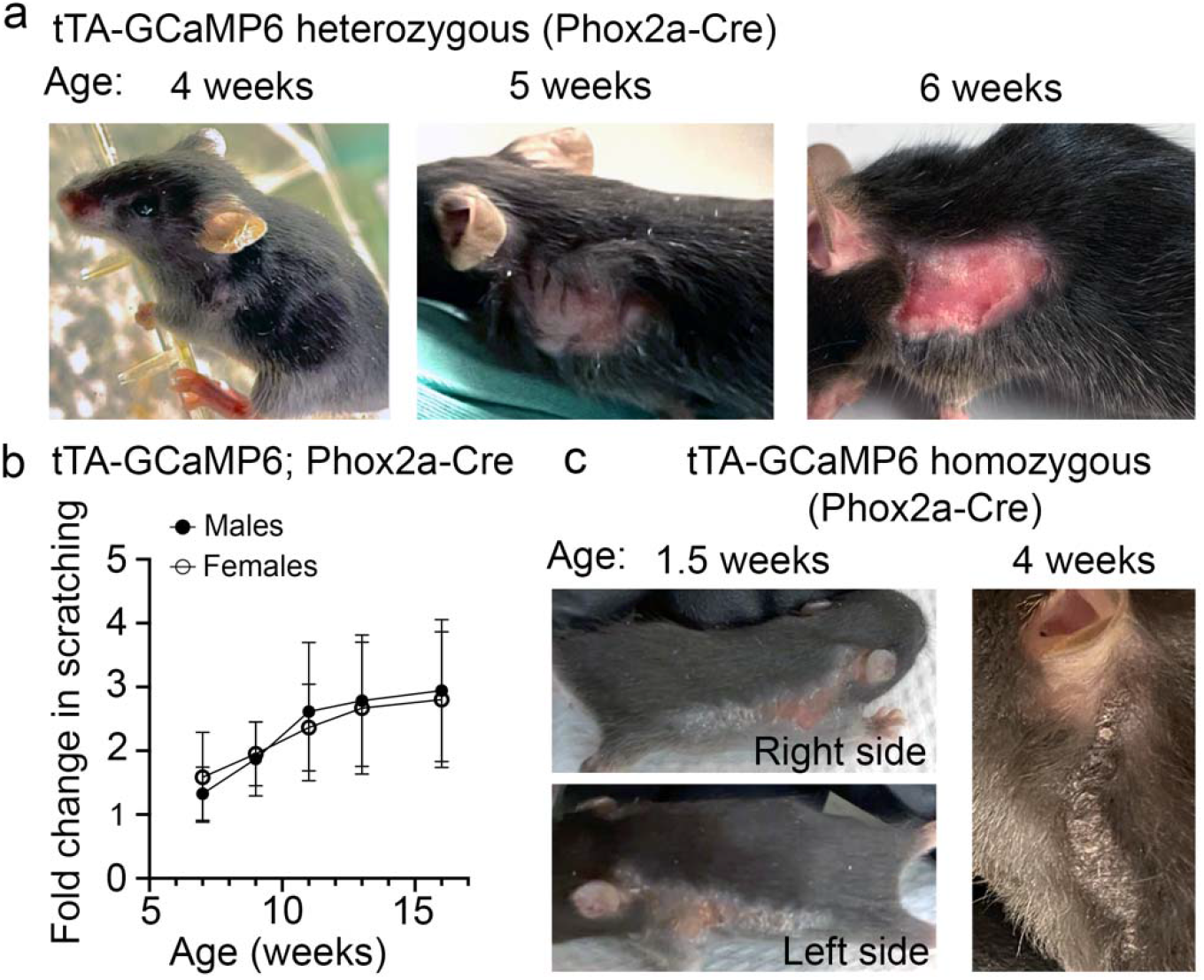
Gene dosage and sex effects on the skin and scratching phenotype. (a) Example of lesion development over time in a Phox2a; tTA-GCaMP6 animal. (b) Male and Female Phox2a; tTA-GCaMP6 mice displayed a similar progression of scratching over time. Data are normalized to the first measurement of scratching at 5 weeks of age. Females (N=6), Males (N=7). (c) Doubling the allele of tTA-GCaMP6 accelerates skin pathology of the shoulder area.

**Supplemental Figure 2:**
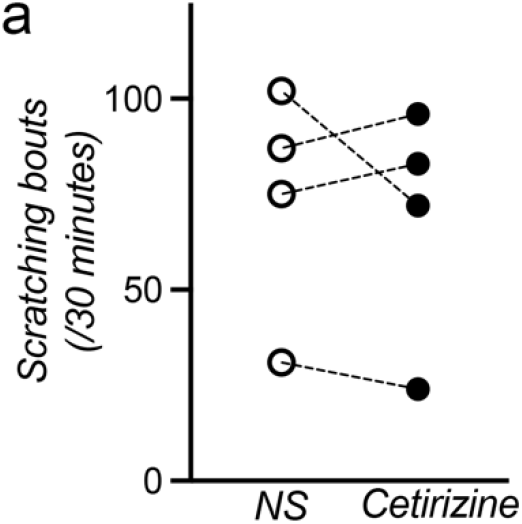
Antihistamines do not reduce spontaneous scratching in Phox2a; tTA-GCaMP6 mice. (a) No difference in scratching bouts after ip. injection of the antihistamine, Cetirizine. NS = normal saline control

**Supplemental Figure 3:**
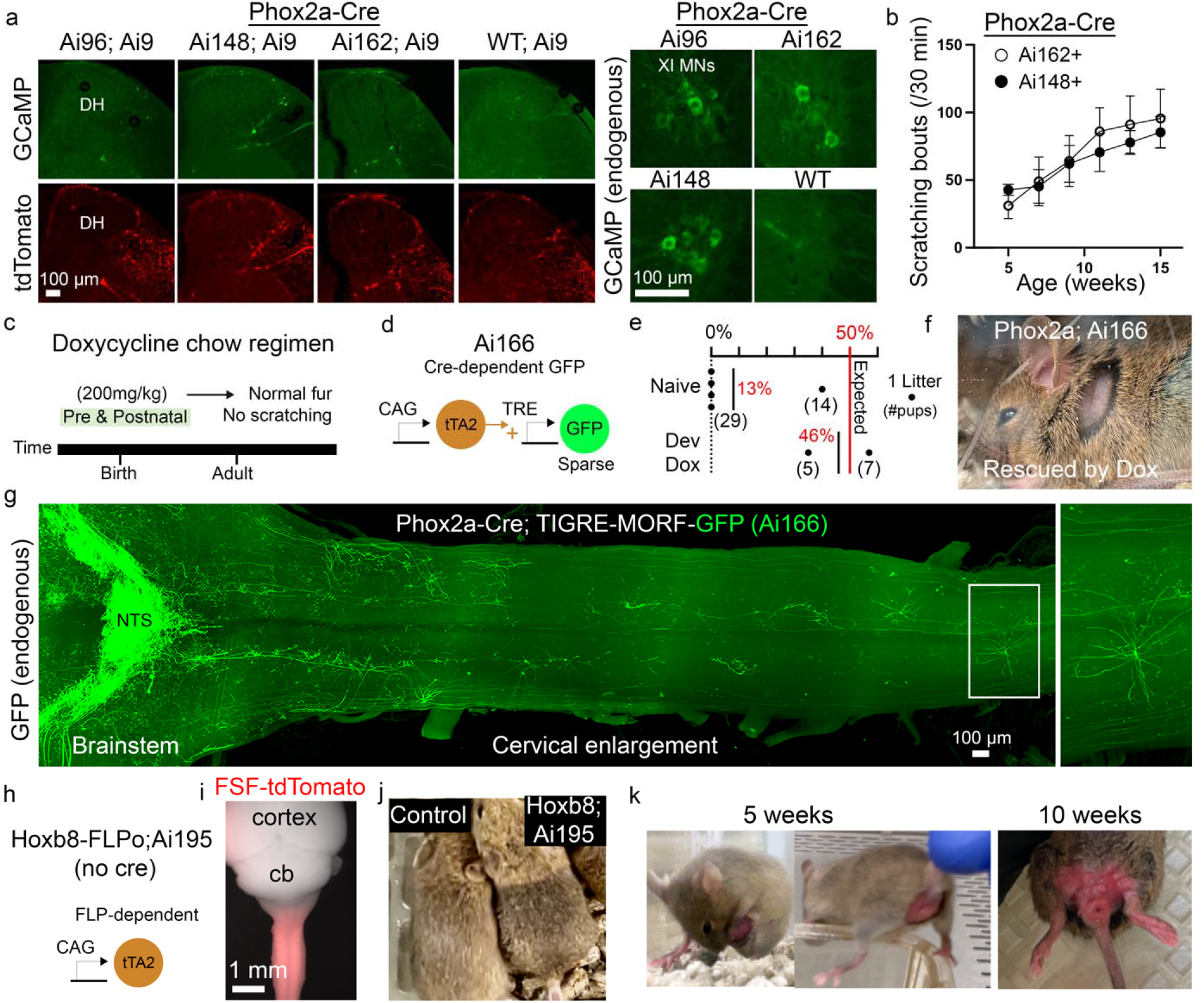
CAG-tTA2 overexpression without GCaMP6 leads to additional pathologies. (a) Histological comparison of endogenous GCaMP fluorescence in mice of the indicated genotypes: cervical dorsal horn neurons (Left) and cervical XI motoneuron populations (Right). tdTomato+ neurons in all genotypes verify recombination activity. (b) No significant difference in the time course of scratching between the fast and slow variants of GCaMP6. Ai162 (N=9), Ai148 (N=4). (c) Scheme of doxycycline chow administration to inhibit tTA activity. Dox given during pre and postnatal periods prevented scratching-induced lesions. (d) The TIGRE2.0 variant, Ai166, does not contain GCaMP6. (e) The percentage yield of viable double transgenic (Phox2a-Cre and Ai166) mice per litter is plotted in two conditions: Naïve and Developmental Dox. The expected yield of both conditions (50%) is noted by a red line compared to the actual % yield of each group noted in red text. (f) Image of a Dox-rescued Phox2a-Cre; Ai166 mouse at 5 weeks of age. (g) Histology of a rescued Phox2a-Cre; Ai166 mouse. The spinal cord was cleared using the SHIELD method (Park et al. 2018) (h-k) The TIGRE2.0 variant, Ai195, expresses tTA2 after FLP recombination. When Ai195 is crossed to Hoxb8-FLPo mice that display selective recombination caudal to C7, coat darkening, and repeated biting/licking behavior are provoked in double transgenic mice. Skin lesions develop in this genotype around the genital area, rather than the shoulder, and increase in size over time.

**Supplemental Figure 4:**
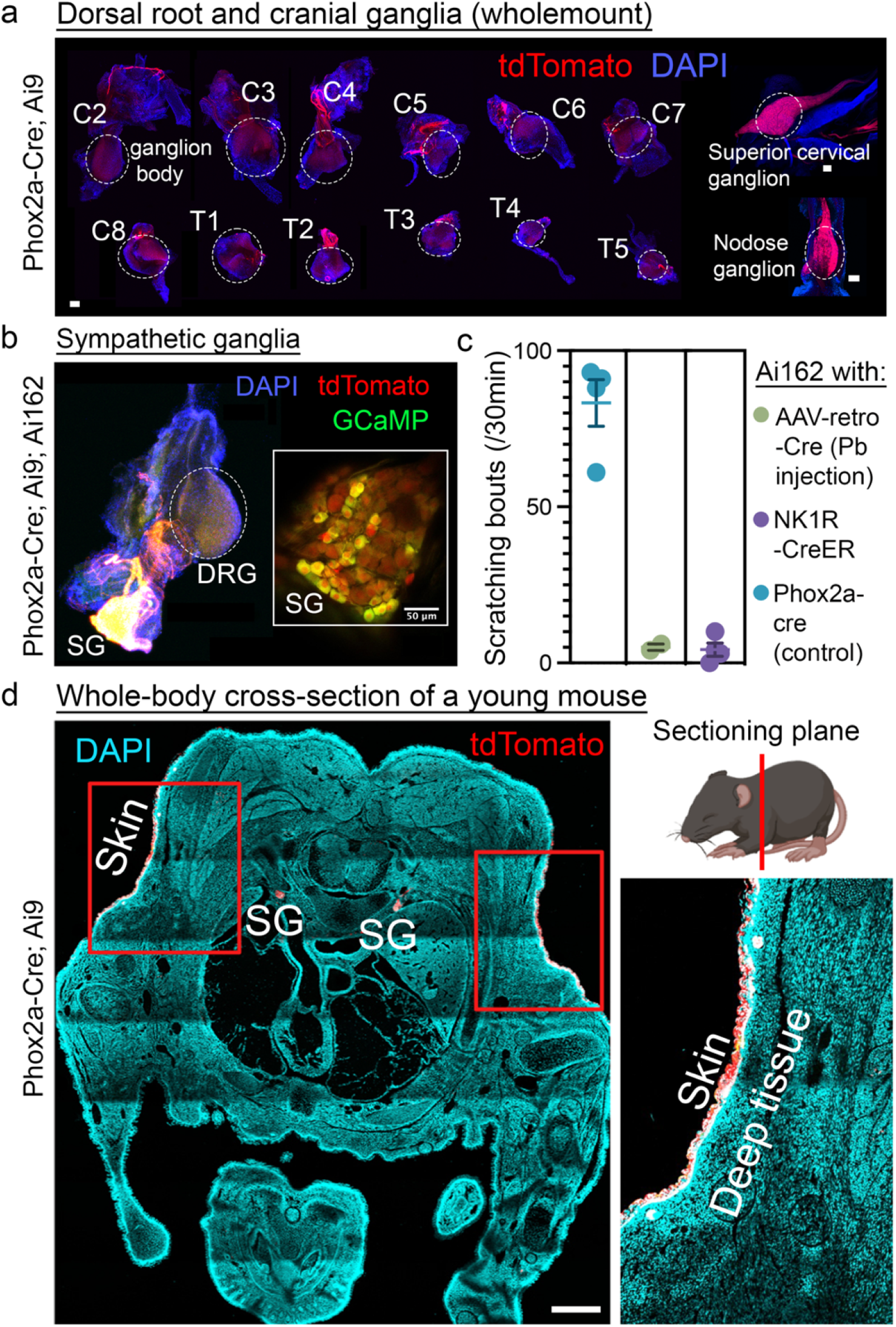
Additional Phox2a-Cre lineage fate mapping. (a) Fate-mapped tdTomato+ neurons are not found in the DRGs. Cervical DRG levels are shown and annotated. In contrast to DRGs, cranial vagal ganglia are fully labeled with tdTomato+ neurons (right). Scale bars = 200 µm. (b) A sympathetic ganglion (SG) attached to the T10 DRG expresses GCaMP and tdTomato. (c) Neither NK1R-CreER nor retrograde strategies to activate tTA in dorsal horn projection neurons induced scratching. (d) Cross-section from a P7 mouse reveals prominent tdTomato reporter expression in the epidermis and SG at the shoulder level. Scale bar = 1 mm.

**Supplemental Figure 5:**
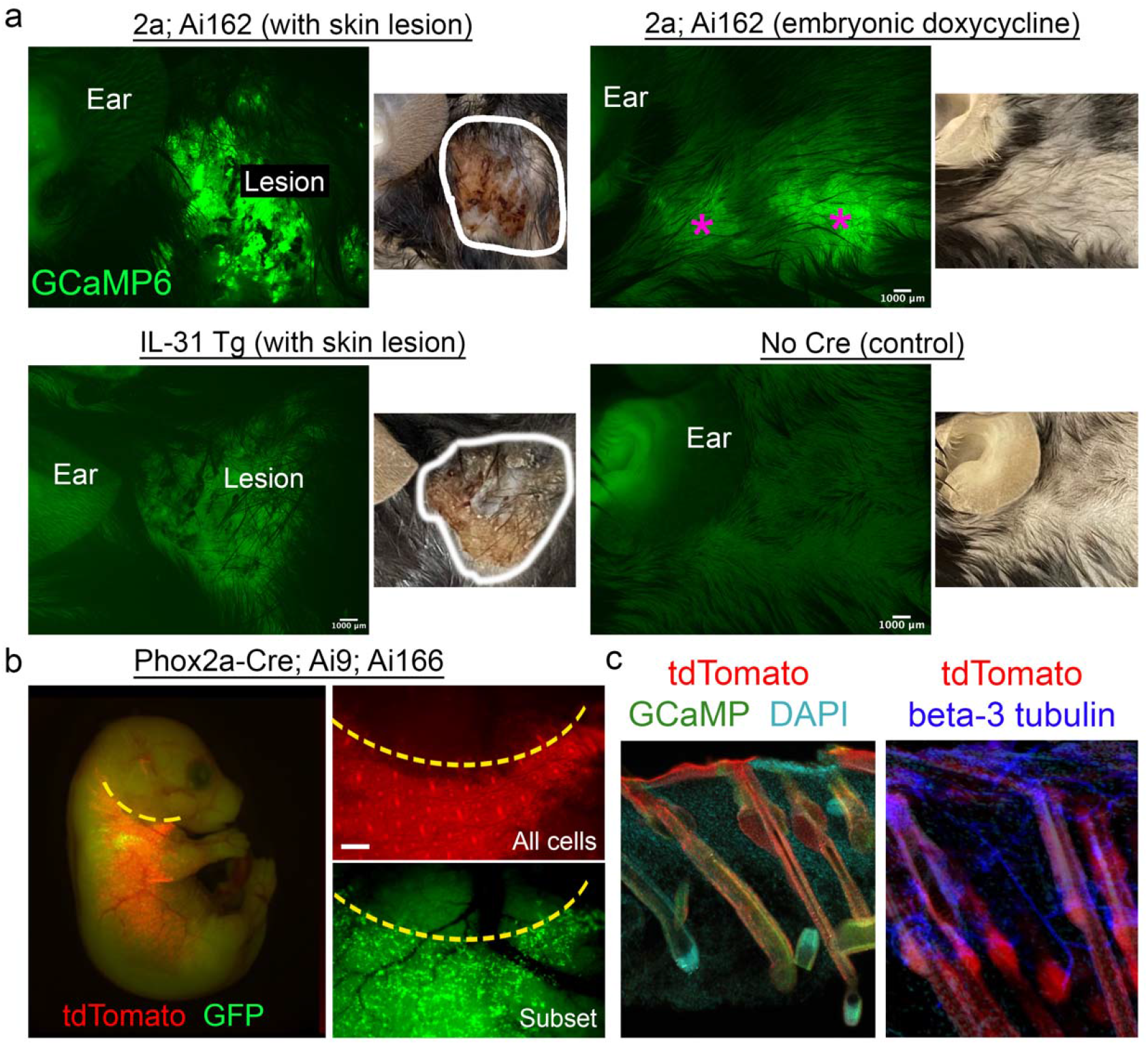
Additional profiling of Phox2a-keratinocytes. (a) Wide field of view image of GCaMP6 fluorescence illustrating skin lesions in various genotypes and conditions. Lesions (white circles of real color images) only show bright signals in a Phox2a-Cre; Ai162 mouse. In a Phox2a; Ai162 mouse without lesions, due to Dox administration during development, GCaMP+ keratinocytes are found in patches (purple asterisks) scattered throughout the shoulder skin (upper right). No labeling is found in tTA-GCaMP6 mice without Cre (lower right) (b) A Phox2a; Ai166; Ai9 mouse illustrates the entire lineage of Phox2a-keratinocytes in red and a random subset of the lineage in green. Scale bar = 100 µm. (c) Hair follicles of triple transgenic mice illustrate GCaMP and tdTomato signal from resident keratinocytes. Beta-3 tubulin+ sensory neuron terminals localize near the fate-mapped keratinocytes of the skin and hair follicles.

## References

1. Ahanonu, Biafra, Andrew Crowther, Artur Kania, Mariela Rosa Casillas, and Allan Basbaum. 2023. “Long-Term Optical Imaging of the Spinal Cord in Awake, Behaving Animals.” BioRxiv: The Preprint Server for Biology, May. https://doi.org/10.1101/2023.05.22.541477.

2. Akiyama, Tasuku, Tony Nguyen, Eric Curtis, Katsuko Nishida, Jahnavi Devireddy, Jeremy Delahanty, Mirela Iodi Carstens, and Earl Carstens. 2015. “A Central Role for Spinal Dorsal Horn Neurons That Express Neurokinin-1 Receptors in Chronic Itch.” Pain 156 (7): 1240–46.

3. Ballantyne, Jane C., Alan B. Loach, and Daniel B. Carr. 1988. “Itching after Epidural and Spinal Opiates.” Pain 33 (2): 149–60.

4. Baumbauer, Kyle M., Jennifer J. DeBerry, Peter C. Adelman, Richard H. Miller, Junichi Hachisuka, Kuan Hsien Lee, Sarah E. Ross, H. Richard Koerber, Brian M. Davis, and Kathryn M. Albers. 2015. “Keratinocytes Can Modulate and Directly Initiate Nociceptive Responses.” ELife 4 (September). https://doi.org/10.7554/eLife.09674.

5. Berger, S. L., W. D. Cress, A. Cress, S. J. Triezenberg, and L. Guarente. 1990. “Selective Inhibition of Activated but Not Basal Transcription by the Acidic Activation Domain of VP16: Evidence for Transcriptional Adaptors.” Cell 61 (7): 1199–1208.

6. Bohic, Manon, Aman Upadhyay, Jaclyn T. Eisdorfer, Jessica Keating, Rhiana Simon, Brandy Briones, Chloe Azadegan, et al. 2023. “Developmentally Determined Intersectional Genetic Strategies to Dissect Adult Sensorimotor Function.” BioRxiv. https://doi.org/10.1101/2022.05.16.492127.

7. Braz, Joao M., Todd Dembo, Alexandra Charruyer, Ruby Ghadially, Marlys S. Fassett, and Allan I. Basbaum. 2021. “Genetic Priming of Sensory Neurons in Mice That Overexpress PAR2 Enhances Allergen Responsiveness.” Proceedings of the National Academy of Sciences of the United States of America 118 (8). https://doi.org/10.1073/pnas.2021386118.

8. Brunet, Jean-François, and Alexandre Pattyn. 2002. “Phox2 Genes - from Patterning to Connectivity.” Current Opinion in Genetics & Development 12 (4): 435–40.

9. Daigle, Tanya L., Linda Madisen, Travis A. Hage, Matthew T. Valley, Ulf Knoblich, Rylan S. Larsen, Marc M. Takeno, et al. 2018. “A Suite of Transgenic Driver and Reporter Mouse Lines with Enhanced Brain-Cell-Type Targeting and Functionality.” Cell 174 (2): 465–480.e22.

10. Damke, H., M. Gossen, S. Freundlieb, H. Bujard, and S. L. Schmid. 1995. “Tightly Regulated and Inducible Expression of Dominant Interfering Dynamin Mutant in Stably Transformed HeLa Cells.” Methods in Enzymology 257: 209–20.

11. Frateschi, Simona, Eric Camerer, Giovanna Crisante, Sarah Rieser, Mathieu Membrez, Roch-Philippe Charles, Friedrich Beermann, et al. 2011. “PAR2 Absence Completely Rescues Inflammation and Ichthyosis Caused by Altered CAP1/Prss8 Expression in Mouse Skin.” Nature Communications 2 (January): 161.

12. Fritzsch, Bernd, Karen L. Elliott, and Joel C. Glover. 2017. “Gaskell Revisited: New Insights into Spinal Autonomics Necessitate a Revised Motor Neuron Nomenclature.” Cell and Tissue Research 370 (2): 195–209.

13. Gill, G., and M. Ptashne. 1988. “Negative Effect of the Transcriptional Activator GAL4.” Nature 334 (6184): 721–24.

14. Gong, Peng, Matthew J. Epton, Guoliang Fu, Sarah Scaife, Alexandra Hiscox, Kirsty C. Condon, George C. Condon, et al. 2005. “A Dominant Lethal Genetic System for Autocidal Control of the Mediterranean Fruitfly.” Nature Biotechnology 23 (4): 453–56.

15. Gossen, M., and H. Bujard. 1992. “Tight Control of Gene Expression in Mammalian Cells by Tetracycline-Responsive Promoters.” Proceedings of the National Academy of Sciences of the United States of America 89 (12): 5547–51.

16. Han, Harry J., Carolyn C. Allen, Christie M. Buchovecky, Michael J. Yetman, Heather A. Born, Miguel A. Marin, Shaefali P. Rodgers, et al. 2012. “Strain Background Influences Neurotoxicity and Behavioral Abnormalities in Mice Expressing the Tetracycline Transactivator.” The Journal of Neuroscience: The Official Journal of the Society for Neuroscience 32 (31): 10574–86.

17. Hirsch, Marie-Rose, Fabien d’Autréaux, Susan M. Dymecki, Jean-François Brunet, and Christo Goridis. 2013. “A Phox2b::FLPo Transgenic Mouse Line Suitable for Intersectional Genetics.” Genesis 51 (7): 506–14.

18. Huang, Huizhen, Marissa S. Kuzirian, Xiaoyun Cai, Lindsey M. Snyder, Jonathan Cohen, Daniel H. Kaplan, and Sarah E. Ross. 2016. “Generation of a NK1R-CreER Knockin Mouse Strain to Study Cells Involved in Neurokinin 1 Receptor Signaling.” Genesis 54 (11): 593–601.

19. Ikoma, Akihiko, Martin Steinhoff, Sonja Ständer, Gil Yosipovitch, and Martin Schmelz. 2006. “The Neurobiology of Itch.” Nature Reviews. Neuroscience 7 (7): 535–47.

20. Jouvet, Nathalie, Khalil Bouyakdan, Scott A. Campbell, Cindy Baldwin, Shannon E. Townsend, Maureen A. Gannon, Vincent Poitout, Thierry Alquier, and Jennifer L. Estall. 2021. “The Tetracycline-Controlled Transactivator (Tet-On/Off) System in β-Cells Reduces Insulin Expression and Secretion in Mice.” Diabetes 70 (12): 2850–59.

21. Kukreja, L., R. Shahidehpour, G. Kim, J. Keegan, K. R. Sadleir, T. Russell, J. Csernansky, et al. 2018. “Differential Neurotoxicity Related to Tetracycline Transactivator and TDP-43 Expression in Conditional TDP-43 Mouse Model of Frontotemporal Lobar Degeneration.” The Journal of Neuroscience: The Official Journal of the Society for Neuroscience 38 (27): 6045–62.

22. Liu, Boyi, Yan Tai, Satyanarayana Achanta, Melanie M. Kaelberer, Ana I. Caceres, Xiaomei Shao, Jianqiao Fang, and Sven-Eric Jordt. 2016. “IL-33/ST2 Signaling Excites Sensory Neurons and Mediates Itch Response in a Mouse Model of Poison Ivy Contact Allergy.” Proceedings of the National Academy of Sciences of the United States of America 113 (47): E7572–79.

23. Lozano-Gerona, Javier, Aida Oliván-Viguera, Pablo Delgado-Wicke, Vikrant Singh, Brandon M. Brown, Elena Tapia-Casellas, Esther Pueyo, et al. 2020. “Conditional KCa3.1-Transgene Induction in Murine Skin Produces Pruritic Eczematous Dermatitis with Severe Epidermal Hyperplasia and Hyperkeratosis.” PloS One 15 (3): e0222619.

24. McKinney, B. C., J. S. Schneider, G. L. Schafer, J. L. Lowing, S. Mohan, M. X. Zhao, M. Y. Heng, et al. 2008. “Decreased Locomotor Activity in Mice Expressing TTA under Control of the CaMKII Alpha Promoter.” Genes, Brain, and Behavior 7 (2): 203–13.

25. Mikesell, Alexander R., Olena Isaeva, Francie Moehring, Katelyn E. Sadler, Anthony D. Menzel, and Cheryl L. Stucky. 2022. “Keratinocyte PIEZO1 Modulates Cutaneous Mechanosensation.” ELife 11 (September). https://doi.org/10.7554/eLife.65987.

26. Moehring, Francie, Ashley M. Cowie, Anthony D. Menzel, Andy D. Weyer, Michael Grzybowski, Thiago Arzua, Aron M. Geurts, Oleg Palygin, and Cheryl L. Stucky. 2018. “Keratinocytes Mediate Innocuous and Noxious Touch via ATP-P2X4 Signaling.” ELife 7 (January). https://doi.org/10.7554/eLife.31684.

27. Mohan, Hemanth, Xu An, X. Hermione Xu, Hideki Kondo, Shengli Zhao, Katherine S. Matho, Bor-Shuen Wang, Simon Musall, Partha Mitra, and Z. Josh Huang. 2023. “Cortical Glutamatergic Projection Neuron Types Contribute to Distinct Functional Subnetworks.” Nature Neuroscience 26 (3): 481–94.

28. Nguyen, Eileen, Grace Lim, Huiping Ding, Junichi Hachisuka, Mei-Chuan Ko, and Sarah E. Ross. 2021. “Morphine Acts on Spinal Dynorphin Neurons to Cause Itch through Disinhibition.” Science Translational Medicine 13 (579). https://doi.org/10.1126/scitranslmed.abc3774.

29. Ottina, Eleonora, Victor Peperzak, Katia Schoeler, Emma Carrington, Roswitha Sgonc, Marc Pellegrini, Simon Preston, Marco J. Herold, Andreas Strasser, and Andreas Villunger. 2017. “DNA-Binding of the Tet-Transactivator Curtails Antigen-Induced Lymphocyte Activation in Mice.” Nature Communications 8 (1): 1028.

30. Park, Young-Gyun, Chang Ho Sohn, Ritchie Chen, Margaret McCue, Dae Hee Yun, Gabrielle T. Drummond, Taeyun Ku, et al. 2018. “Protection of Tissue Physicochemical Properties Using Polyfunctional Crosslinkers.” Nature Biotechnology, December. https://doi.org/10.1038/nbt.4281.

31. Pattyn, A., X. Morin, H. Cremer, C. Goridis, and J. F. Brunet. 1997. “Expression and Interactions of the Two Closely Related Homeobox Genes Phox2a and Phox2b during Neurogenesis.” Development 124 (20): 4065–75.

32. Pedro, P., N. Salinas Parra, and R. Iglesias-Bartolome. 2019. “349 Regulation of Epithelial Stem Cell Proliferation and Differentiation by the Gαi Heterotrimeric G Protein.” The Journal of Investigative Dermatology 139 (5): S60.

33. Roome, R. Brian, Farin B. Bourojeni, Bishakha Mona, Shima Rastegar-Pouyani, Raphael Blain, Annie Dumouchel, Charleen Salesse, et al. 2020. “Phox2a Defines a Developmental Origin of the Anterolateral System in Mice and Humans.” Cell Reports 33 (8): 108425.

34. Rossi, Jari, Nina Balthasar, David Olson, Michael Scott, Eric Berglund, Charlotte E. Lee, Michelle J. Choi, Danielle Lauzon, Bradford B. Lowell, and Joel K. Elmquist. 2011. “Melanocortin-4 Receptors Expressed by Cholinergic Neurons Regulate Energy Balance and Glucose Homeostasis.” Cell Metabolism 13 (2): 195–204.

35. Salghetti, S. E., A. A. Caudy, J. G. Chenoweth, and W. P. Tansey. 2001. “Regulation of Transcriptional Activation Domain Function by Ubiquitin.” Science 293 (5535): 1651–53.

36. Shimada, Steven G., and Robert H. LaMotte. 2008. “Behavioral Differentiation between Itch and Pain in Mouse.” Pain 139 (3): 681–87.

37. Stanke, M., D. Junghans, M. Geissen, C. Goridis, U. Ernsberger, and H. Rohrer. 1999. “The Phox2 Homeodomain Proteins Are Sufficient to Promote the Development of Sympathetic Neurons.” Development 126 (18): 4087–94.

38. Steinmetz, Nicholas A., Christina Buetfering, Jerome Lecoq, Christian R. Lee, Andrew J. Peters, Elina A. K. Jacobs, Philip Coen, et al. 2017. “Aberrant Cortical Activity in Multiple GCaMP6-Expressing Transgenic Mouse Lines.” ENeuro 4 (5). https://doi.org/10.1523/ENEURO.0207-17.2017.

39. Tiveron, M. C., M. R. Hirsch, and J. F. Brunet. 1996. “The Expression Pattern of the Transcription Factor Phox2 Delineates Synaptic Pathways of the Autonomic Nervous System.” The Journal of Neuroscience: The Official Journal of the Society for Neuroscience 16 (23): 7649–60.

40. Trier, Anna M., Madison R. Mack, Avery Fredman, Masato Tamari, Aaron M. Ver Heul, Yonghui Zhao, Changxiong J. Guo, et al. 2022. “IL-33 Signaling in Sensory Neurons Promotes Dry Skin Itch.” The Journal of Allergy and Clinical Immunology 149 (4): 1473–1480.e6.

41. Wercberger, Racheli, and Allan I. Basbaum. 2019. “Spinal Cord Projection Neurons: A Superficial, and Also Deep, Analysis.” Current Opinion in Physiology 11 (October): 109– 15.

42. Wilson, Sarah R., Lydia Thé, Lyn M. Batia, Katherine Beattie, George E. Katibah, Shannan P. McClain, Maurizio Pellegrino, Daniel M. Estandian, and Diana M. Bautista. 2013. “The Epithelial Cell-Derived Atopic Dermatitis Cytokine TSLP Activates Neurons to Induce Itch.” Cell 155 (2): 285–95.

43. Yang, Yaxiong, Nan Liu, Yuanyuan He, Yuxia Liu, Lin Ge, Linzhi Zou, Sen Song, Wei Xiong, and Xiaodong Liu. 2018. “Improved Calcium Sensor GCaMP-X Overcomes the Calcium Channel Perturbations Induced by the Calmodulin in GCaMP.” Nature Communications 9 (1): 1504.

44. Veldman, Matthew B., Chang Sin Park, Charles M. Eyermann, Jason Y. Zhang, Elizabeth Zuniga-Sanchez, Arlene A. Hirano, Tanya L. Daigle, et al. 2020. “Brainwide Genetic Sparse Cell Labeling to Illuminate the Morphology of Neurons and Glia with Cre-Dependent MORF Mice.” Neuron 108 (1): 111–27.e6.

45. Cevikbas, Ferda, Joao M. Braz, Xidao Wang, Carlos Solorzano, Mathias Sulk, Timo Buhl, Martin Steinhoff, and Allan I. Basbaum. 2017. “Synergistic Antipruritic Effects of Gamma Aminobutyric Acid A and B Agonists in a Mouse Model of Atopic Dermatitis.” The Journal of Allergy and Clinical Immunology 140 (2): 454–64.e2.

